# Evaluation of UMAP as an alternative to t-SNE for single-cell data

**DOI:** 10.1101/298430

**Authors:** Etienne Becht, Charles-Antoine Dutertre, Immanuel W. H. Kwok, Lai Guan Ng, Florent Ginhoux, Evan W. Newell

**Affiliations:** Singapore Immunology Network (SigN), Agency for Science, Technology and Research (A*STAR)

## Abstract

Uniform Manifold Approximation and Projection (UMAP) is a recently-published non-linear dimensionality reduction technique. Another such algorithm, t-SNE, has been the default method for such task in the past years. Herein we comment on the usefulness of UMAP high-dimensional cytometry and single-cell RNA sequencing, notably highlighting faster runtime and consistency, meaningful organization of cell clusters and preservation of continuums in UMAP compared to t-SNE.

## Introduction

The last decades have witnessed a large increment in the number of parameters analysed in single cell cytometry studies. It currently reaches around 20 for flow-cytometry, 40 for mass-cytometry, and more than 20,000 in single-cell RNA-sequencing. In this context, dimensionality reduction techniques have been pivotal in enabling researchers to visualize high-dimensional data. While principal component analysis has historically been the main technique used for dimensionality reduction (DR), the recent years have highlighted the importance of non-linear DR techniques to avoid overcrowding issues. Common such techniques[1] include Isomap, Diffusion Map and t-SNE[2] (also renamed viSNE[3]). t-SNE is currently the most commonly-used technique and is efficient at highlighting local structure in the data, which for cytometry notably translates to the representation of cell populations as distinct clusters. t-SNE however suffers from limitations such as loss of large-scale information (the inter-cluster relationships), slow computation time and inability to meaningfully represent very large datasets[4]. A new algorithm, called Uniform Manifold Approximation and Projection (UMAP) has been recently published by McInnes and Healy[5]. They claim that compared to t-SNE it preserves as much of the local and more of the global data structure, with a shorter runtime. Since t-SNE has been extremely prevalent in the field of cytometry broadly encompassing flow and mass-cytometry as well as single-cell RNA-sequencing (scRNAseq), we tested these claims on well-characterized single-cell datasets[6-8]. We confirm that UMAP is an order of magnitude faster than t-SNE. In addition to this straightforward advantage, we argue that UMAP is not only able to create informative clusters, but is also able to organize these clusters in a meaningful way. We illustrate these claims by showing that UMAP can order clusters from T and NK cells from 8 human organs[7] in a way that both identifies major cell lineages but also a common axis that broadly recapitulates their differentiation stages. We also show that UMAP allows for an easier visualization of multibranched cellular trajectories by using a mass-cytometry[6] and a scRNAseq[8] datasets both recapitulating hematopoiesis.

## Results

### Faster runtime, equivalent local information and superior global structure

We ran UMAP and t-SNE simultaneously on a dataset covering 39 samples originating from 8 distinct human tissues enriched for T and NK cells, of more than >350,000 events with 42 protein targets[7]. As observed by McInnes and Healy[5], we measured runtimes that were significantly lower (5 minutes on average for UMAP for 200,000 cells, versus 2 hour and 22 minutes for Barnes-Hut t-SNE) across a large range of dataset sizes (Figure S1). Using Phenograph[9] clustering and manual cluster labeling, we classified events into 7 broad cell populations (Figure S2A). UMAP and t-SNE were both successful at pulling together only clusters corresponding to similar cell populations with generally very good correspondence with Phenograph clustering (Figure 1A and Figure S2B). However t-SNE separated cell populations into distinct clusters more commonly than UMAP, notably splitting CD8 T cells, gamma-delta T cells and contaminating cells (likely including B cells) into two distinct clusters each. Although this highlights that tSNE might be more sensitive in segregating these populations that differ, we were unable to test this quantitatively. We also note that although these cells were not always segregated into completely distinct clusters by UMAP, these cell populations remained similarly identifiable in UMAP as compared to tSNE (Figure S2B). In addition, UMAP appeared more stable than t-SNE, being more consistent across distinct replicates and independent subsampling which should facilitate consistency in its intepretation (Figure S3). By color-coding the tissues of origin on the UMAP and t-SNE maps, we observed that t-SNE grouped cell clusters according to their origin more often than UMAP (Figure 1B and Figure S4). UMAP instead ordered events according to their origin within each major cluster, roughly from cord-blood and PBMCs, to liver and spleen, and to tonsils one the one end to skin, gut and lung on the other end. The sample type was not given as an input of any of these two algorithms. Instead we observed that UMAP was able to recapitulate the differentiation stage of T cells within each major cluster, as seen by the expression levels of events for the resident-memory markers CD69 and CD103, the memory T cell marker CD45RO and naive cells marker CCR7 on the UMAP projection (Figure 1C). By contrast, while t-SNE identified similar continuums within clusters, they had no apparent structure along a common axis that made them easily identifiable (Figure 1D).

**Figure 1.**
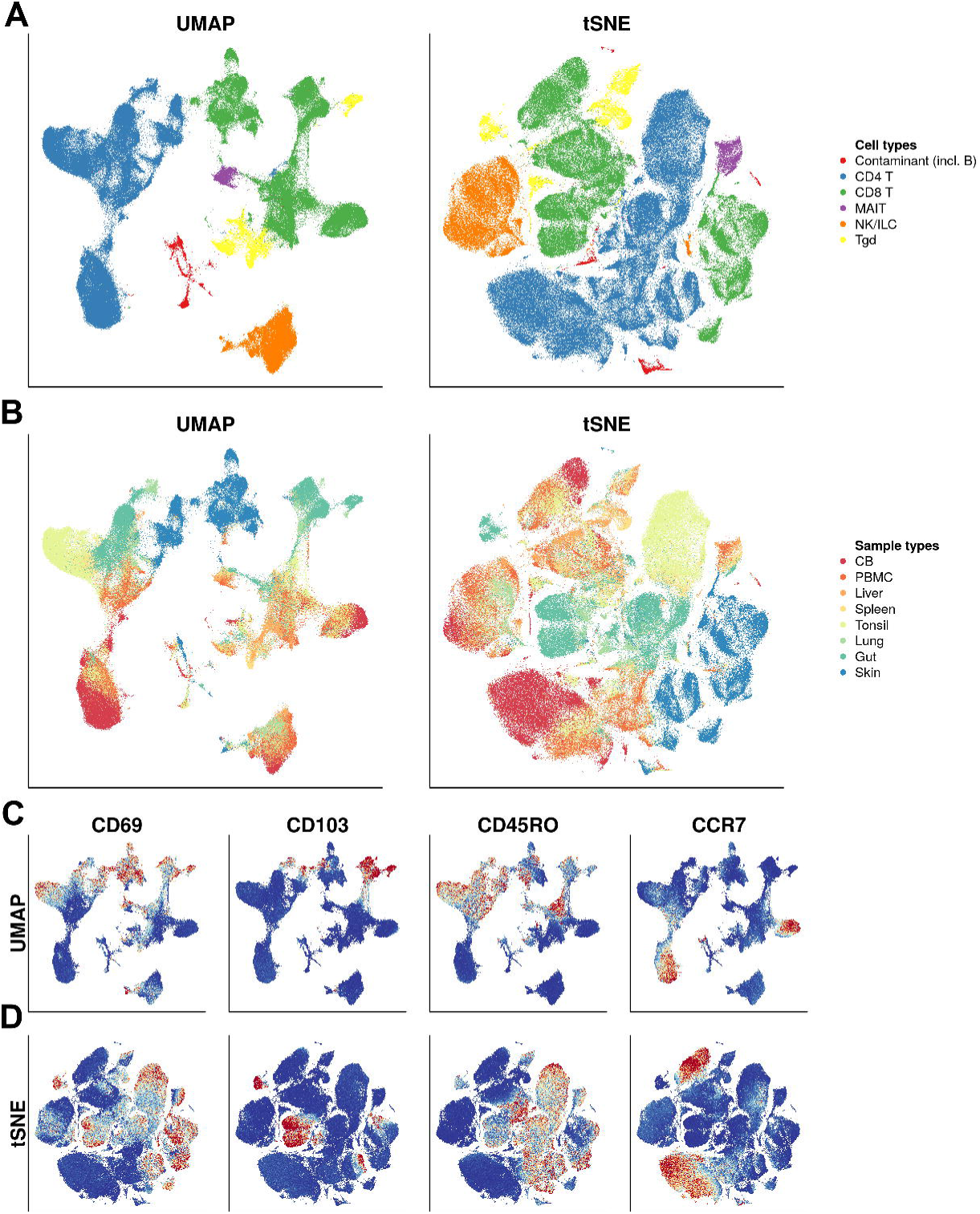
Experimental design and work flow. UMAP and t-SNE projections of the Wong et al. dataset colored according to **A)** broad cell lineages, **B)** tissue of origin, and for **C)** UMAP and **D)** t-SNE, the expression of CD69, CD103, CD45RO and CCR7. For **C)** and **D)**, blue denotes minimal expression, beige intermediate and red high expression.

### UMAP better represents the multi-branched trajectory of hematopoietic development

To investigate how UMAP handles continuity of cell phenotypes we applied it alongside t-SNE on the well-documented topic of bone-marrow hematopoiesis using both a mass-cytometry (>86,000 events, 25 parameters, 24 cell populations annotated by its authors[6]) and a scRNAseq dataset (three sample types, 51,252 cells, 25,912 dimensions[8]). On the mass-cytometry dataset, UMAP visually revealed 8 major cell clusters (Figure 2A). One was composed of all B cell subsets (and close to a small cluster of plasma cells) and one of all T cell subsets. Four small homogeneous clusters corresponded to macrophages, NK cells, eosinophils and non-classical monocytes. The last cluster contained 11 out of the 24 manually-gated populations and appeared most interesting with respect to hematopoiesis. Indeed, these populations were ordered according to a five-leaf branched structure that was consistent with hematopoietic differentiation: hematopoietic stem cells (HSC) overlapped with multipotent progenitors (MPP). These cells neighbored common lymphoid progenitors (CLP) on one side, and common myeloid progenitors (CMP) on the other. CMP led to myeloid-erythroid progenitors (MEP) which led to unlabelled erythrocytes (Figure S5), and to granulocyte-myeloid progenitors (GMP). GMP then led to classical monocytes that further led to myeloid dendritic cells on one branch and to cells labeled as intermediate monocytes on another branch. UMAP linked basophils to a population of Lin^−^cKit^+^Sca1^−^CD34^+^FcγRII/III^+^FcεRIα^+^ cells, consistently with a previously-described phenotype for committed basophil progenitors[10]. These putative progenitors appeared closer to CMP than GMP, a topic that is still intensely debated. Neutrophils were gated-out from the dataset by its authors and are thus absent from this representation[6]. t-SNE identified relatively similar clusters with a few differences (Figure 2B), notably singling out more clusters than UMAP. CD4^+^ T cells were separated from other T cell subsets. As noted by others[11], t-SNE expands low density areas and tends to ignore global relationships. Thus, while some paths from HSC and MPP to differentiated populations were still apparent-notably from HSC to monocytes, the overall structure was less clear, as no narrow “neck” led to larger terminal clusters. t-SNE also separated basophils from their putative precursors close to CMP and GMP and pDC from CLP. The density of events in the dimensionally-reduced space also appeared less uniform in t-SNE, with large clusters in the t-SNE space being less dense than the smaller ones. In contrast, the density of UMAP clusters appeared more uniform, which could help avoid biases in interpreting phenotypic heterogeneity in large versus small clusters (Figure S6).

**Figure 2.**
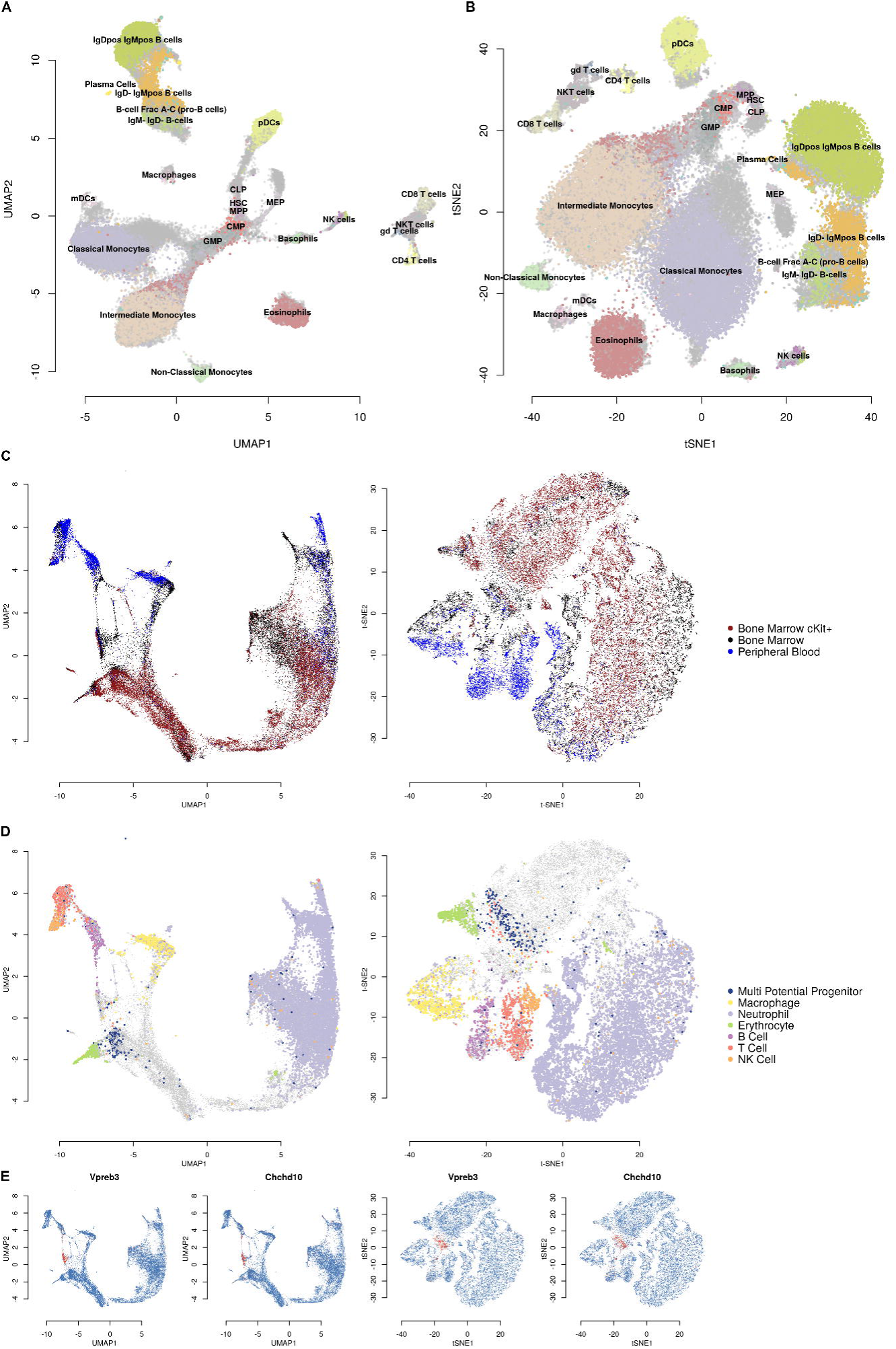
**A)** UMAP and **B)** t-SNE projection of the Samusik_01 dataset. Events are color-coded according to manual gates provided by the authors of the dataset. **C)** UMAP and t-SNE projections of the Han dataset, color-coded by tissue of origin or **D)** by cell populations. **E)** Expression of the *V-set pre-B cell surrogate light chain 3* gene (Vpreb3) and *Chchd10* genes on the UMAP and t-SNE projections of the Han dataset. Blue denotes minimal expression, beige intermediate and red high expression.

From the scRNAseq dataset we analyzed collectively the transcriptomes of cells isolated from Bone Marrow (BM), cKit^+^ BM and Peripheral Blood (PB) to facilitate identifying mature versus progenitor cell populations (Figure 2C). We first removed low-abundance cell types such as basophils and eosinophils, contaminants such as mature erythrocytes as well as outlier cells originating from unique samples and highly expressing mitochondrial transcripts (Figure S7). Using published cell signatures specific for mouse BM cell populations[12], we were able to identify cell clusters that corresponded to MPP, MEP, macrophages, B cells, T cells and NK cells (Figure 2D). Consistently with the UMAP projection of the mass-cytometry dataset, MPP were found in the middle of a larger group of clusters that led to differentiated cells originating from PB samples (Figure 2C). PB events consisted of distinct clusters of lymphocytes (T, NK and B cells), macrophages, MEP and neutrophils (Figure 2D). Although this does not prove that cells lying between MPPs and differentiated cells are committed progenitors, these results suggest that UMAP could be used as a hypothesis generating tool to identify putative markers for such cells. By investigating a small cluster of cells lying in between MPP and mature B cells in the UMAP projection, we were indeed able to identify the pre-B cell marker Vpreb3[13] and to hypothesize that Chchd10 could be another gene marker for pre-B cells in mouse bone marrow (Figure 2E). These conclusions and hypotheses would have been more difficult to draw using t-SNE which blurred the relationship of terminal clusters to MPPs (Figure 2D and Figure 2E).

## Discussion

Our analysis and example provided show that UMAP seems to yield representation that are as meaningful as t-SNE does, particularly in its ability resolve even subtly differing cell populations. In addition, it provides the useful and intuitively pleasing feature that it preserves more of the global structure, and notably, the continuity of the cell subsets. In addition to making plots easier to interpret, we highlight that this also improves its utility for generating hypotheses related to cellular development. On a practical level, UMAP outputs are faster to compute and more reproducible than those from t-SNE. Altogether, based on its ease of use, these results and our other experience so far, we anticipate that UMAP will be a highly valuable tool that can be rapidly adopted by single-cell analysis community.

## Methods

### Datasets

The main characteristics of the datasets we analyzed are presented in Table 1. For the Wong et al. dataset, based on live CD45+ cell events available on FlowRepository (see table) non-B (CD19^-^), non-monocyte (CD14^-^) were selected using FlowJo software. In order to partially equalize weighting of each human tissue, a maximum of 10,000 events were randomly sampled from each of the 39 samples prior to analysis. Other datasets were used as described in the table.

| Dataset | Identifier | Single-cell<br>technique | Organism | Tissues | Samples | Single<br>cells | Analyzed<br>parameters |
| --- | --- | --- | --- | --- | --- | --- | --- |
| Wong | FlowRepository<br>FR-FCM-ZZTM | Mass-cytometry | Human | 8 distinct<br>tissues | 39 | 327,457 | 39 |
| Samusik_01 | FlowRepository<br>FR-FCM-ZZPH | Mass-cytometry | Mouse | Bone<br>marrow | 1 | 86,864 | 38 |
| Han | Figshare<br>865e694ad06d5<br>857db4b | Single-cell<br>RNAseq | Mouse | Bone<br>marrow<br>and blood | 14 | 51,252 | 25,912 |

### Transformations and pre-processing

For the bone marrow mass-cytometry data we used an arcsinh transformation with a cofactor of 1, and a logicle transform (parameters w=0.25, t=16409, m=4.5, a=0) for the Wong dataset. For the scRNAseq dataset we transformed count into reads per millions (thus normalizing the number of counts per cells to 1).

### Running UMAP and t-SNE

For both mass-cytometry datasets we used UMAP using 15 nn, a min_dist of 0.2 and euclidean distance. For the scRNAseq dataset we computed 100 approximate principal components using the *IRLBA* R package and used them as an input for both t-SNE and UMAP. We then ran UMAP using 30 nearest neighbors (nn) and a *min_dist* of 0.1 and using the “*correlation*” metric. For t-SNE we ran the Barnes-Hut[14] implementation of the t-SNE algorithm through its R implementation in the *Rtsne* package, using default parameters.

### Cell annotations

For the Samusik_01 dataset we used cell annotations provided by the authors and available from the public repository. For the Wong et al. dataset we used Phenograph clustering (with default parameters k=30) and manually labeled the clusters into broad cell populations. For then Han et al. dataset we used the AUCell R package[15], which computes the AUC of gene sets within each single cell, using gene sets from the Haemopedia[12] resource to annotate cell lineages. We then manually thresholded these AUC scores to obtain categorical labels. Cells that were assigned to multiple lineages were set to unlabeled.

---

*Supplementary Figure 1*

Runtime of both UMAP (red) and t-SNE (blue) on randomly-selected subsets of the Wong dataset using various sampling sizes. 3 subsamples were selected for each subset size and input to both algorithms. Vertical lines represent standard deviations - and are too short to be visible for most data points.

*Supplementary Figure 2*

**A)** Phenotypic characterization of the phenograph clusters. Each cluster medoid is represented after column-wise Z-score transformation. **B)** Identification of each phenograph cluster of both UMAP (left) and t-SNE. For clarity, only twelve clusters are shown per plot.

*Supplementary Figure 3*

Datapoints were colored according to their position on the UMAP (left) or t-SNE (right) projection for the full Wong dataset. Then 3 subsets of various sizes were randomly selected and input to UMAP and t-SNE. The resulting projections were colored according to the full dataset projections in order to compare positions across random subsets and replicates.

*Supplementary Figure 4*

UMAP and t-SNE projections of the Wong dataset individually color-coded by tissue of origin.

*Supplementary Figure 5*

Expression of Ter119 (a marker for mature erythrocytes) on the UMAP projection of the Samusik_01 dataset.

*Supplementary Figure 6*

Heatmap of the density of a 300×300 square grid of the UMAP or t-SNE projections for the Samusik_01 dataset. The number of events in each bin is color-coded.

*Supplementary Figure 7*

Top: UMAP projection of the full Han dataset annotated by AUC scores for various cell lineages (red: high score, blue: low score). Bottom: full Han dataset colored by sample type, Sample ID and pre-filtering status.

## Acknowledgements

We thank members of the Singapore Immunology Network and notably members of the E.N. laboratory. We thank Shamin Li, Yannick Simoni, Melissa Chng, Yang Cheng, Jack Wee Lim and Michael Fehlings for their insightful feedbacks. This study was funded by A-STAR/SIgN core funding and A-STAR/SIgN immunomonitoring platform funding.

